# Discordant evolution of organellar genomes in peas (*Pisum* L.)

**DOI:** 10.1101/2020.05.19.104224

**Authors:** Vera S. Bogdanova, Natalia V. Shatskaya, Anatoliy V. Mglinets, Oleg E. Kosterin, Gennadiy V. Vasiliev

## Abstract

Plastids and mitochondria have their own small genomes which do not undergo meiotic recombination and may have evolutionary fate different from each other and nuclear genome, thus highlighting interesting phenomena in plant evolution. We for the first time sequenced mitochondrial genomes of pea (*Pisum* L.), in 38 accessions mostly representing diverse wild germplasm from all over pea geographical range. Six structural types of pea mitochondrial genome were revealed. From the same accessions, plastid genomes were sequenced. Bayesian phylogenetic trees based on the plastid and mitochondrial genomes were compared. The topologies of these trees were highly discordant implying not less than six events of hybridisation of diverged wild peas in the past, with plastids and mitochondria differently inherited by the descendants. Such discordant inheritance of organelles is supposed to have been driven by plastid-nuclear incompatibility, known to be widespread in pea wide crosses and apparently shaping the organellar phylogenies. The topology of a phylogenetic tree based on the nucleotide sequence of a nuclear gene *His5* coding for a histone H1 subtype corresponds to the current taxonomy and resembles that based on the plastid genome. Wild peas (*Pisum sativum* subsp. *elatius* s.l.) inhabiting Southern Europe were shown to be of hybrid origin resulting from crosses of peas similar to those presently inhabiting south-east and north-east Mediterranean in broad sense.

## 1. Introduction

The genus *Pisum* L. (Fabaceae, Fabea) includes an ancient (Zohary & Hopf, 2000) but still important crop, the pea (*Pisum sativum* L.), and its wild relatives (Maxted & Ambrose, 2001), a potential source of genetic diversity for crop improvement. Besides, wild peas are a good model for studying microevolution being annual plants with very small populations. At the same time wild peas are still insufficiently studied and their diversity was obscured by a number of circumstances: (i) taxonomic instability of the genus (Maxted & Ambrose, 2001; Coulot & Rabaute, 2016; 2017), (ii) confusion of genuine wild plants with ‘primitive’, traditionally cultivated local landraces or feral cultivated peas; (iii) spontaneous crossing while reproduction in germplasm collections; (iv) confusion of labels and physical contamination of seeds in germplasm collections; (v) explicit or hidden duplication of accessions in different collections (Kosterin 2016; Zaytseva et al., 2017), (vi) depletion and extinction of wild pea populations due to deterioration of natural habitats (Maxted and Kell, 2009). These complications, especially (i) and (ii), eluded the interpretation of the results of repeated phylogenetic and diversity analyses (Ellis et al., 1998; Vershinin et al., 2003; Jing et al., 2005; 2007; 2010; Tar’an et al., 2005; Smýkal et al., 2017). While the methods applied were powerful in resolving clusters and evolutionary branches, their proper interpretation demanded a clear idea of the taxonomical treatments of accessions involved, their cultivation status, phenotype, ecology and geography.

Here we adopt the taxonomic system of *Pisum* outlined by Maxted & Ambrose (2001). According to this generalised system, the genus embraces a clear-cut wild species *P. fulvum* Sibth. et Smith from Israel, Syria, SE Turkey and Sinai Peninsula and a diverse species *P. sativum* containing both wild (conventionally considered as the subspecies *P. sativum* subsp. *elatius* (Bieb.) Aschers. et Graebn. in a broad sense) and cultivated (*P. sativum* subsp. *sativum*) peas. Besides, there is a cultivated taxon from Yemen and Ethiopia, resembling *P. sativum* but scarcely reproductively compatible with it, of an equivocal rank (Kosterin, 2017; Weeden, 2018), here considered as a species *P. abyssinicum*.

The key feature of wild peas is dehiscing pods, providing ballistic seed dispersal (Weeden, 2007; Ambrose and Ellis, 2008; Zaytseva et al., 2017). Plants with non-dehiscent pods cannot persist in nature, while plants with dehiscing pods cannot be harvested. This exerts a situation of disruptive selection forcing wild peas to remain wild, cultivated peas to remain cultivated, and products of rare crosses between them to join either of the two gene pools with respect to pod dehiscence. Another important adaptation of wild peas to their unstable habitats is seed dormancy, absent in cultivated peas (Weeden, 2007).

We have compiled a reliable collection of about 60 accessions of truly wild peas with dehiscing pods and reliable labels, which, together with a cultivated pea collection, allows to investigate patterns of the genus divergence. First we worked out a simple diagnostic system of three dimorphic molecular markers, *rbcL, cox1* and *SCA*, from the plastid, mitochondrial and nuclear genomes, respectively, allowing to classify overwhelming majority of peas into three major groups with respect to three most frequent allelic combinations of those markers. These combinations were denoted as A (*rbcL* with the restriction site for *Hsp* AI, *cox1* with the restriction site for *Psi* I, fast SCA electromorph), C (differing from A by the absence of the *Psi* I recognition site in *cox1*) and B (the mentioned restriction sites missing both from *rbcL* and *cox1*, slow *SCA* electromorph) (Kosterin and Bogdanova, 2008; Kosterin et al., 2010). From this point of view we attempted reconstruction of phylogeography of wild peas (Kosterin et al., 2010; Zaytseva et al., 2017). We made a phylogeny reconstruction based on the genes *His5* and *His7* of histone H1 subtypes 5 and 7, of which the former appeared a good phylogenetic marker (Zaytseva et al., 2012; 2015). In the resulting trees, based on just two concatenated nuclear genes (Zaytseva et al., 2015), three main clades were resolved: (1) including only *P. fulvum*, with combination A; (2) with only wild representatives of *P. sativum*, with combinations A and C (‘clade AC’), and (3) containing wild and cultivated accessions of *P. sativum*, with combination B (‘clade B’) (some peas with rare marker combination appearing in the two latter). The cultivated subspecies *P. sativum* subsp. *sativum* formed a branch inside the lineage B, while *P. abyssinicum* (with combination A) formed a small branch inside the lineage AC, in accordance with previous phylogenetic studies in the genus (Ellis et al., 1998; Vershinin et al., 2003; Jing et al., 2010; Kreplak et al., 2019), that argues against its species rank (Weeden, 2018). At the same time, carriers of combinations A and C were interspersed in the ‘lineage AC’, that is, the difference between combination C and A consisting in the absence/presence of a recognition site for the *Psi* I endonuclease in the mitochondrial *cox1* gene did not look a phylogenetic landmark.

Since the mentioned evolutionary lineages were isolated on the basis of three markers, of which one was from the plastid genome and one from the mitochondrial genome, it was natural to conduct phylogenetic analyses based on the entire organellar genomes, which contain a great amount of phylogenetically valuable information (Van der Paer et al., 2017) and, unlike the nuclear sequences, have the advantage of avoiding meiotic recombination between parental genomes.

Earlier we undertook sequencing and phylogenetic analysis of the plastid genomes for 22 mostly wild pea accessions (Bogdanova et al., 2018). The reconstructed phylogeny in general corresponded to that obtained on the basis of nuclear histone H1 genes (Zaytseva et al. 2012; 2015) again showing ‘lineages AC and B’, but in addition revealed two unexpected basic divegrences. Accessions of *P. fulvum* clustered with two wild pea accessions classified as *P. sativum* subsp. *elatius* s.l., from Georgia and Turkey. The latter, an enigmatic wild pea accession P 013 from Tokat Province, is remarkable for exhibiting amphicarpy, thought to be confined to *P. fulvum*, that phenotypically proves relevance of its reconstructed phylogenetic position, suggesting its being a ‘missing link’ between *P. sativum* and *P. fulvum* representing an early branching of their common ancestor (Bogdanova et al., 2018). This accession also formed a basic branch on the tree based on a histone H1 gene, *His5* (Zaytseva et al., 2012). The fact that one of diverse wild pea accessions from the Caucasus also belonged to the same branch in the plastid tree was puzzling. The other unexpected branch on the plastid genome tree, the first divergence of the main *P. sativum* clade, was formed by sole accession JI 1096, one of a series of wild pea accessions from the Athos Peninsula, Greece, which looked as typical wild peas of Southern Europe. Hence JI 1096 looked like carrying some cryptic ancient plastid lineage on the otherwise ‘common’ genetic background (Bogdanova et al., 2018).

In the plastid genome tree (Bogdanova et al., 2018), as well as on the histone H1 trees (Zaytseva et al., 2012; 2015), carriers of different alleles of the mitochondrial *cox1* gene did not cluster to each other. Thus it was interesting to reconstruct phylogeny based on the mitochondrial genomes. Until present, no mitochondrial genome of pea has been published. We undertook high throughput sequencing of both organellar genomes in a set of 38 pea accessions, 35 of which represent genuine wild peas, proved to have dehiscing pods. Each accession was represented by a progeny of a single plant randomly chosen from those provided by germpasm collections, or in some cases collected in nature. We assume that a single individual well represents a local wild pea population, since their populations are isolated, very small in number (tens to hundreds of plants) and area occupied (Abbo et al., 2008; 2013; Smýkal et al., 2018; Kosterin et al., 2020) and are characterised by scarce intra-population genetic diversity (Zaytseva et al., 2015; Smýkal et al., 2018).

We compared phylogenetic reconstructions on the basis of mitochondrial genomes and plastid genomes and revealed several principal discordances between them. This result suggests an important role of ancient hybridisation and nuclear-plastid incompatibility in shaping microevolution of peas, and perhaps of many other plants.

## 2. Materials and methods

### 2.1 Plant material

This study is based on 38 pea accessions of mostly wild and some cultivated peas (Table 1) of broad geographical origin and representing a wide range of variation. They were obtained from Agricultural Research Service of the United States Department of Agriculture thanks to the courtesy of Clarice Coyne, from John Innes Centre thanks to the courtesy of Michael Ambrose, from Cornell University thanks to Norman Weeden, from Vavilov All-Union Institute of Plant Breeding, kindly provided by Petr Smykal, or collected from nature by ourselves. In each case, a single plant was chosen and its progeny was involved into analysis.

**Table 1.** Pea accession studied (ordered by provenance from west to east) with accompanying information (in square brackets where reconstructed rather than provided by sources), their constitution of organellar genomes with respect to their main evolutionary branches (see ‘Results’) and Gene Bank accession number of corresponding DNA sequences.

| accession designation adopted in this paper | other known designations/ taxonomic attributions of this accession | origin; date of collection (where relevant/available), collector/originator; reference (if published in a paper) | latitude (N) | longitude (E) | organellar constitution, allele combination of three markers (where available) | ID numbers in databases for plastid genome, mitochondrial genome and *His5* gene (sequences obtained anew underlined) |
| --- | --- | --- | --- | --- | --- | --- |
| *Pisum fulvum*, wild |  |  |  |  |  |  |
| WL 2140 | WL 2029; JI 224; PI 560061 | Israel, Valley of the Cross. | [31.77] | [35.20] | P1 M1, A | MG458702, FR848865, mitogenome ID pending |
| 706 | JI 3268, PI 560065, L96 | Israel, Ruhama, a coastal plain, on calcareous sandstone, in dwarf-shrub formation (Ben-Ze'ev & Zohary, 1973) | [31.5] | [34.7] | P1 M1, A | MG458703, FR848856, mitogenome ID pending |
| PI 343955 | PIG 49, 22645, G 19532 | Turkey, mountains above Kaş (2 km by road), arid limestone E slope, 60 m a.s.l., Coll. May 1969 by H.S. Gentry. (Heterogeneous for very strong vs nil leaflet serration; the non-serrate type analysed) | [36.2] | [29.7] | P1 M1 | pending |
| *Pisum abyssinicum*, cultivated |  |  |  |  |  |  |
| VIR 2759 | WL 1491, WL 2042 | Ethiopia | no data | no data | P4 M2, A | MG859923, FR848852, mitogenome ID pending |
| WL 1446 |  | no data | - “ - | - “ - | P4 M2, A | FR848845, IDs for organellar genomes pending |
| *Pisum sativum* L. subsp. *elatius* (Bieb.) Aschers. et Graebn. s.l., wild |  |  |  |  |  |  |
| IG 64350 |  | Algeria, Blida | [36.5] | [2.8] | P4 M1, A | MG882488, MH269702, mitogenome ID pending |
| CE11 | JI 3557, W6 56891, PE1 | Portugal, Trás-os-Montes e Alto Douro Province, Bragança District, Vimioso Municipality, 1.4 km NE of Uva village, 8.07.2010, O. Kosterin (Zaytseva et al., 2015). | 41.50 | - 6.48 | P4 M3, C | MG882484, LN812960, mitogenome ID pending |
| JI 2724 |  | Spain, Menorca, Mahon | [39.87] | [4.30] | P4 M2, A | FR848867, IDs for organellar genomes pending |
| 723 | JI 3271,<br>PI 560060,<br>L106 | Italy, Sardinia, Cagliari env., in hedges (Ben-Ze'ev & Zohary; 1973). Perhaps a hybrid with *P. sativum* subsp. *elatius* (Zaytseva et al., 2018) | [39.2] | [9.1] | P4 M2, A | FR848846, IDs for organellar genomes pending |
| PIS 2845 |  | Italy, Sicily, Palermo [Prov.], Balze di Cugliace – Palazzo Adriano | [37.7] | [13.4] | P4 M3, C | MG833029, MH269699, mitogenome ID pending |
| PI 344537 |  | Italy, Sicily, Palermo [Prov.], Bosco della Ficuzza. Coll. by A. Di Martino | [37.9] | [13.4] | P4 M4, C | MG882489, FR848890, mitogenome ID pending |
| PIS 2853 |  | Hungary, Baranya, [Siklósiközség, Nagyarsány env.,] Szársomlyó [Mt.]. | [35.85] | [18.41] | P4 M3, C | MG882487, MH269700, mitogenome ID pending |
| JI 1091 | PI 344005,<br>W6 8705,<br>22611 | Greece, Athos Peninsula, Karyes, 400 m a.s.l., among open secondary shrubs, heavy ground litter, upper edge of olive grove. Coll. June 1969 by H.S. Gentry | 40.25 | 24.33 | P4 M4, C | MG859920, FR848866, mitogenome ID pending |
| JI 1096 | PI 344013,<br>W6 8710,<br>22735 | Greece, Athos Peninsula, above Karyes, 450 m a.s.l., chestnut and oak woodland. Coll. June 1969 by H.S. Gentry | 40.25 | 24.33 | P3 M3, C | MG859921, FR848874, mitogenome ID pending |
| W6 10925 |  | Bulgaria, 30 m from beach at Albena, off sandy side of trail, 5 m a.s.l. Coll. 28.06.1992 by R.M. Hanno and W.J. Kaiser | [43.37] | [28.08] | P5 M4, B | pending |
| PI 343974 | 22632,<br>G 23875 | Turkey, [Muğa II], 2 km SE of Selimiye [(Marmaris)] on road to Milas, field margin along stream bank, in *Rubus* thickets, 70 m a.s.l. Coll. May 1969 by H.S. Gentry | [36.68] | [28.12] | P4 M2 | pending |
| P 017 | PI 560973,<br>W6 9813,<br>270685-0105 | Turkey, Mersin II, 19 km of road from Aydinchik to Gulnar, 790 m a.s.l. Coll. 27.06.1985 by W.J. Kaiser, F.J. Muehlbauer (Muehlbauer et al., 1990) | [36.28] | [33.39] | P5 M4, B | FR848843, IDs for organellar genomes pending |
| PI 343999 | 22699 | Turkey, Mersin II, on the road 6 km N of Gülek, scarce in a wheat field, 120 m a.s.l. Coll. May 1969 by H.S. Gentry | [37.30] | [34.77] | P4 M1 | MT468418, IDs for organellar genomes pending |
| Pe 013 | 190785-0105, claimed to be the same as PI 560969 and W6 9809 but these are some different genotype, see (Bogdanova et al., 2018) | Turkey, Tokat Province, the left side of the road from Sivas to Tokat about 5 km from Tokat, 1 km to Geryaz village, a rocky stony slope near water canal, 750 m a.s.l. (Muehlbauer et al., 1990; see comments in Bogdanova et al., 2018) | [40.26] | [36.55] | P2 M2, A | MK440218, FR848882, mitogenome ID pending |
| Pse 001 | 010689-0201, W6 1972 | Turkey, Elazig Il, NW side of Hazar Lake, near cemetery, ca 3km S of Firat University Research Station; on the road from Elazig to Diyarbakyr, 1270 m a.s.l. Coll. 01.06.1989 by W.J. Kaiser, F.J. Muehlbauer, C.R. Sperling. (Muehlbauer et al., 1990) | 38.50 | 39.37 | P3 M4 | MK440219, MT468417, ID for mitochondrial genome pending |
| P 015 | PI 560971, W6 9811, 090785-03 | Turkey, Diyarbakyr Il, on road to Siverek, Karaçadag (Karyarallari Park), 42 km W of Diyarbakyr. Coll. 09.07.1985 by W.J. Kaiser, F.J. Muehlbauer (Muehlbauer et al., 1990) | [37.9] | [39.7] | P5 M4, B | FR848854, IDs for organellar genomes pending |
| Psh 004 | W6 2044, 070689-0202 | Turkey, Mardin Il, 8 km NE of Kiziltepe then 7.7 km NW on road through Kocalar Ortakoy village, 770 m a.s.l., infrequently in terraces, rock fences, ravine slopes on limestone soil. Coll. 07.06.1989 by W.J. Kaiser, F.J. Muehlbauer (Muehlbauer et al., 1990) | 37.30 | 40.63 | P5 M4 | MK455838, IDs for organellar genomes pending |
| Psh 005 | W6 2060, 080689-0403; *P. sativum* subsp. *elatius* var. *pumilio* | Turkey, Mardin Il, 10.7 km S of Midyat on the Midyat-Gizre road. In grassy areas of chickpea field margin; 1050 m a.s.l. Coll. 08.06.1989 by W.J. Kaiser, F.J. Muehlbauer, C.V. Sperling (Muehlbauer et al., 1990) | 37.37 | 41.42 | P5 M4 | pending |
| JI 1794 | 716, 'northern *humile*' | Golan Heights, ca. 3 km NW of Quneitra, Tell Abu Nida (Ben-Ze'ev & Zohary; 1973). | [33.1] | [35.8] | P5 M4, B | HG966675, FR846218, mitogenome ID pending |
| 721 | JI 3262, PI 560058, L104, *P. elatius* | Israel, Mt. Carmel, 5 km NE of Zikhron Ya'akov, in maquis (Ben-Ze'ev & Zohary, 1973). | [32.59] | [35.00] | P4 M2, A | HG966673, FR848849, mitogenome ID pending |
| 712 | JI 3273, PI 560069, L100, 'southern *humile*' | Israel, 10 km S of Be'er Sheva; wadi bed with loess deposit, as a weed in barley (Ben-Ze'ev & Zohary, 1973) | [31.15] | [34.80] | P4 M1, A | HG966676, FR848840, mitogenome ID pending |
| JI 3233 |  | Syria | no data | no data | P5 M4, B | MG882485, MH269701, mitogenome ID pending |
| VIR 320 |  | probably a hybrid of a wild pea from Palestine and the cultivated pea (Bogdanova et al., 2018) | - | - | P4 M1, A | HG966672, FR848838, mitogenome ID pending |
| CE1 | JI 2629, W6 56882 | USSR, Crimea, Simeiz, open juniper forest on a coastal slope. Coll. 6.07.1991 by Y. Trusov, I. Makunin and O. Kosterin | 44.40 | 33,98 | P5 M4, B | MG859922, FR848851, mitogenome ID pending |
| CE3 | W6 56884 | Russia, Krasnodarskiy Kray, Anapa Municipality, 1.5 km ESE of Malyy Utrish village., 47 m a.s.l., a dry glade in open oak-juniper-pistachio stand on a SW slope. Coll. 22.07.2017 by O.E. Kosterin, V.S. Bogdanova | 44.7050 | 37.4750 | P5 M4 | MK952764, IDs for organellar genomes pending |
| CE9 | W6 56889 | Iran, Ostan-e Lorestan Ostan, Shakhrestan-e Aligudarz, Bakhsh-e Besharat, 700 m N of the centre of Kagelestan-e Bar Aftab village, 1841 m a.s.l., the Rudarb-e Aligudarz River valley right slope with open stand of *Prunus scoparia* and grazed by cattle. Coll. 23.05.2017 by coll. O. Kosterin (Kosterin et al., 2020) | 33.0369 | 49.6564 | P5 M4 | MK952766, IDs for organellar genomes pending |
| W6 26112 | G1001 | Georgia, [Shida Kartli Province], Ateni Sioni Church, south facing slope, basaltic area with many stones and boulders and some grass. Coll. 1.07.2004 by F.J. Muehlbauer, M. Akhalkatsi, W.J. Kaiser | 41.904 | 44.094 | P5 M4 | pending |
| W6 26109 | G0801 | Georgia, [Kvemo Kartli Region, Trialeti Range], 20 km of Manglisi, 7 km N of Partsklisi, a roadside slope with weeds and some grasses, 278 m a.s.l. Coll. 30.06.2004 by F.J. Muehlbauer, M. Akhalkatsi, W.J. Kaiser | 41.602 | 44.522 | P2 M1, A | MG882486, MH269698, mitogenome ID pending |
| VIR 2998 | *P. elatius* subsp. *caspicum* Govorov | Georgia; received by VIR from Riga Botanical Garden; a subline from a heterogeneous accession | no data | no data | P5 M4, B | FR848886, IDs for organellar genomes pending |
| IG 140562 |  | Azerbaijan, Shamakhi [District] | 40.69 | 48.64 | P5 M4 | MT468420, IDs for organellar genomes pending |
| *Pisum sativum* subsp. *sativum*, cultivated |  |  |  |  |  |  |
| WL 1238 |  | testerline by H. Lamprecht (Anonymus, 1984) | not appl. | not appl. | P5 M3, B | HG966674, FR848836, mitogenome ID pending |
| WL 1072 |  | - “ - | - “ - | - “ - | P5 M4, B | MG917089, MH269697, mitogenome ID pending |
| Cameor |  | cultivar | - “ - | - “ - | P5 M4 | pending |
| *Vavilovia formosa* (outgroup) |  |  |  |  |  |  |
| - |  | Russia, North Osetia (Shatskaya et al., 2019) (organellar genomes analysed) | no data | no data | not appl. | MK604478, MK48602 and MK48603, |
| - |  | Turkey (Zaytseva et al., 2015) (*His5* gene analysed) | - “ - | - “ - | - “ - | HG934385 |

### 2.2. Total and organellar DNA extraction, PCR amplification and DNA sequencing

Genomic DNA was extracted using a protocol described in Bogdanova et al. (2012). Organellar DNA was extracted according to the protocol of plastid DNA extraction by Jansen et al. (2005) with modifications as described in Bogdanova et al. (2015). One step of the protocol, aimed to separate plastids from mitochondria, was omitted, namely, the centrifugation for 17 min at 4 C at 20,000 g in a step gradient consisting of 6 ml 60% suchrose overlayed with 20 ml of 40% suchrose. By this, DNA of both organelles was isolated. This approach was similar to that of Siniauskaya et al. (2020).

PCR amplification and Sanger sequencing of the *His5* gene were made at the SB RAS Genomics Core Facility as described in Bogdanova et al. (2018).

### 2.3. Ion Torrent PGM sequencing and assembly of organellar genomes

Ion Torrent PGM sequencing was performed at IC&G Center of Genomic Investigations as described in Bogdanova et al. (2015).

To assemble mitochondrial genomes we used the same array of data which was used for the assembly of plastid genomes. It was possible since the read pool in about 10% consisted of mitochondrial DNA (Bogdanova et al., 2015). First, the mitochondrial genome of the WL1238 accession was assembled. Assembly was made starting from a complement of contigs obtained in *de novo* assembly without reference sequence by MIRA4 (Chevreux et al., 1999). A contig consisting of mitochondrial DNA was selected by BLAST search (Altshul et al., 1990) at www.ncbi.nlm.nih.gov using mitochondrial DNA of *Vicia faba* L. (KC189947) as a query. Then the initial stretch of mitochondrial DNA was elongated by an overlapping contig which was searched with the Search function of the Tablet 1.13.07.31 assembly viewer (Milne et al., 2013). When more than one contig overlapped with a nascent mitochondrial assembly, the choice was made by chance. The elongation process was made in both directions until the forming large contig closed into a circle or a sigma-like structure. To procced with the assembly, the second starting mitochondrial DNA stretch was chosen on the basis of its homology with *V. faba* mitochondrial genome not yet included in the assembly. This was elongated in a similar way until it closed into a circle or merged with a circle already assembled. The latter was possible due to the contings refuted by the random choice when constructing the first circle. Then the third stretch of mitochondrial DNA not yet included into the assembly was chosen by homology with *V. faba* mitochondrion and the procedure was repeated. The process lasted until there remained regions of *V. faba* mitochondrial DNA homologous to pea sequences and not covered by the assembly constructed. In the case of WL1238 three large circles were obtained with the sizes of about 72, 125, 165 kb, the latter including a stem-loop structure.

Overlaps of these three circles were identified using the blastn algorithm at blast.ncbi.nlm.nih.gov. Four overlaps of about 200 bp were found (schematically presented in Fig. 1). Evidently, there are various manners to resolve the overlapping circular DNAs into a master circle since each join/repeat has two entrances and two exits that implies four possible configurations of the DNA pass, in a manner of the ‘8’ figure or as two circles. This fact probably reflects the nature of plant mitochondrial DNA which exists as a set of interconverting sub-genomic molecules due to homologous recombination between repeated regions (Gualberto et al., 2014; Kozik et al., 2019). In fact, each of the alternative configurations was represented by some contigs of the starting set. The resulting assembly, however, should have included only two of four possible configurations. The length of overlaps was about 200 bp that is comparable with the mean length of an IonTorrent read which is 188 bp. Therefore, to choose the best way of resolving the joins, using the mirabait utility of the MIRA package (Chevreux et al., 1999), we filtered the reads which crossed both boundaries of a join; the number of such reads was 4 to 10 per join. The DNA pass of individual reads was checked. In three cases it was represented by only two configurations that indicated the best way to resolve the joins. In the fourth case, of 10 reads that crossed both boundaries, 3 were of one type and 6 of another supporting the ‘8’ configuration, and 1 of the third type, supporting the ‘two circles’ configuration. This join was resolved as to form configuration supported by the majority of reads. Totally, two of possible joins were resolved in a manner of the figure ‘8’ and two – in a manner of non-overlapping circles. The DNA pass in the resulting master circle is presented in Fig. 1 as dashed line.

**Figure 1.**
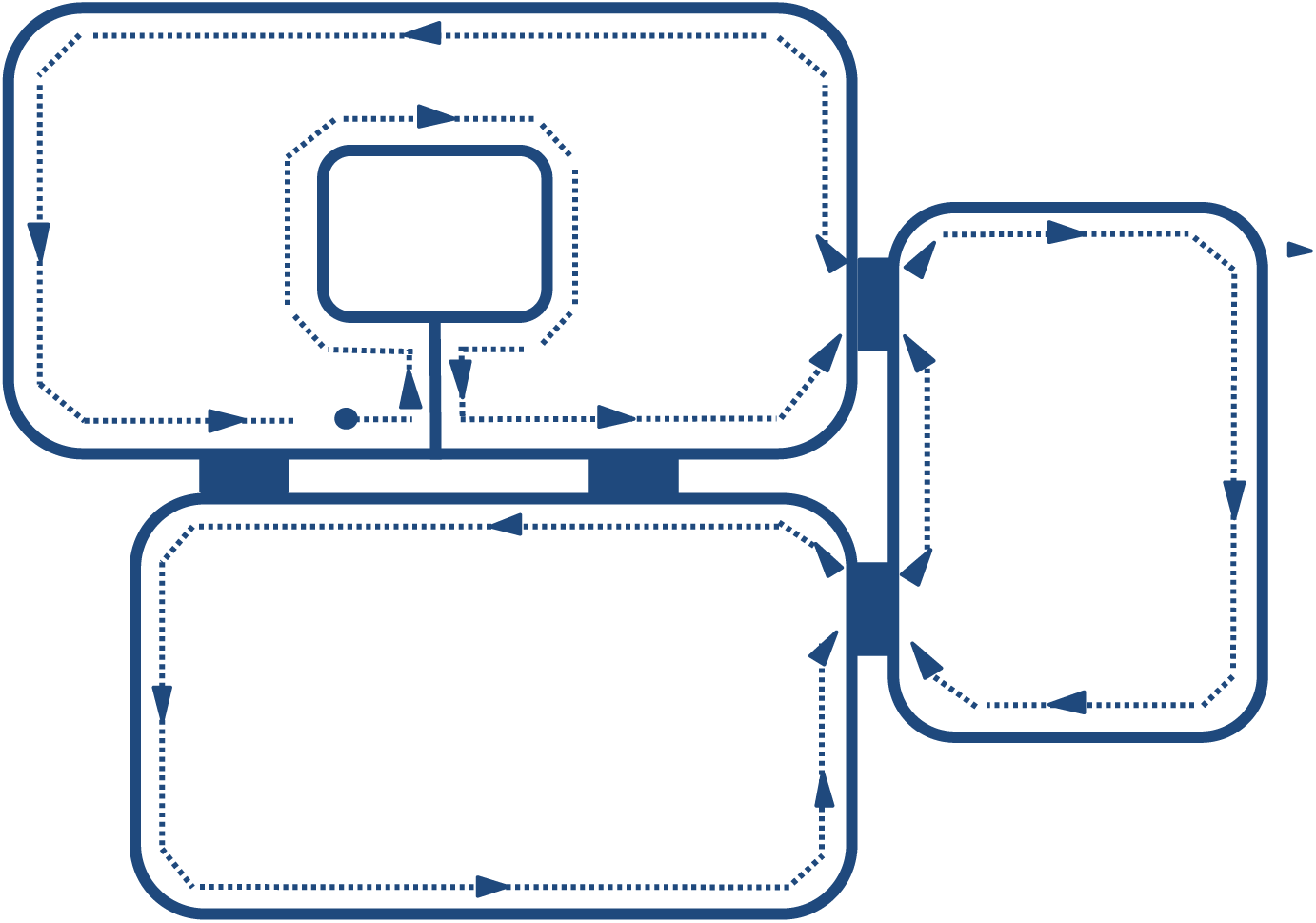
Schematic presentation of mitochondrial genome assembly of the accesssion WL1238. Solid lines represent assembled circular contigs, solid rectangles – overlapping DNA stretches, dashed line with arrowheads represents the DNA pass in the master molecule assembly. The solid circle is the starting point in the resulting assembly.

The reconstructed mitogenome of WL1238 was then used as reference to map mitochondrial genomes of the rest accessions. However, in some pea lines, mitochondrial DNA could not be assembled on the backbone of that of WL1238 and presented obvious cases of rearrangements, as well as large deletions/insertions. In these cases, the entire assembling procedure was made *de novo* as described above.

Plastid genomes were assembled as described in Bogdanova et al. (2015; 2018) using the MIRA4 software (Chevreux et al., 1999).

### 2.4. Alignment and evaluation of nucleotide variation of organellar genomes

The assembled organellar genomes were aligned by ClustalW (Larkin, 2007) program incorporated into Mega 6 package (Tamura et al., 2013) and manually adjusted.

In a number of accessions, the reconstructed mitogenomes had large deletions, insertions, structural rearrangements as compared to the reference mitogenome of WL1238. To align them, the DNA sequence was broken into 5 to 13 parts which were then manually put in an order and orientation corresponding to the reference sequence.

The plastid and mitochondrial genomes were compared to that of WL1238 (*P. sativum* subsp. *sativum*) and the differences registered, are summarized in Appendices A and B, respectively.

### 2.5. Phylogenetic tree construction

Bayesian MCMC analysis was performed with the use of BEAST 2.4.3. software (Drummond & Rambaut, 2007). jModelTest 2.1.10 (Darriba et al., 2012; Guindon and Gascuel, 2003) was used to estimate substitution rates, proportion of invariant sites, Gamma shape parameter and to choose the GTR+I+G model. An uncorrelated lognormal relaxed clock model and Yule process of speciation were applied. One MCMC analysis was run for 100 million generations.

Computations were made at Computational Facility of the Siberian Supercomputer Center SB RAS and Computational Facility of Novosibirsk State University. Trees were visualized in FigTree 1.4.3 (http://tree.bio.ed.ac.uk/software/figtree/ by A.Rambaut).

The reconstructions were rooted with the same outgroup, *Vavilovia formosa* (Stev.) A. Fed. (Table 1), a perennial plant most related to peas (*Pisum* L.). Its plastid and mitochondrial genomes were earlier sequenced by Shatskaya et. al. (2019) and the *His5* gene by Zaytseva et al. (2015). To root a reconstruction based on the mitochondrial genomes, that of *Vicia faba* (KC189947) was also used.

## 3. Results

### 3.1. Size and structure of pea mitochondrial genomes

The lenght of the assembled mitogenomes varied in the range of 346,959– 385,511 bp. According to their length, the assembled mitogenomes fell into three groups. The shortest mitogenomes, 346,959 and 354,692 bp belonged to the abyssinian pea accessions, WL 1446 and VIR 2759, respectively. The most abundant group of 28 accessions had mitogenomes of 363,029–363,928 bp, the third group of 7 accessions had longer mitogenomes – 379,890 to 379,987, and the accession 712 had the longest mitogenome of 385,511 bp due to an about 15 Kbp-repeat, however, in the latter case, assembly artifact is not excluded. The size differences were due to large insertions/deletions. In comparison with the mitogenome of WL1238 taken as reference, longer mitogenomes of the third group differed by about 32 Kb of inserted and about 17 Kb deleted DNA. The shortest mitogenome of the WL1446 abyssinian pea differed by about 17 Kb of deleted DNA, mitogenome of VIR2759 differed from that of WL 1238 by about 15 Kb deleted and about 6 Kb inserted DNA. These large insertions/deletions, however, concerned non-coding regions and did not change the gene content. The only exception was the above mentioned 15-Kb repeat in 712 which contained three tRNA genes.

Apart from the size differences, the assembled mitogenomes differed by the gene order. Although the employed assembly algorithm admits ambiguities, possible variants of the resulting assemblies are compatible with the complements of the original reads. On the contrary, the revealed distinct structural types cannot be assembled from the given read set. Based on the criterion of compatibility with the read sets, six structural types of mitochondrial genomes were distinguished, which corresponded well to the above mentioned size groups. The most abundant 363–364 Kb group included two structural types which could be interconverted by two recombination events. Two other structural types were represented by two shorter *P. abyssinicum* mitogenomes, that of VIR 2759 arranged as one master circle and that of WL 1446 arranged as a sigma-like structure. The other two structural types were observed in the group of longer genomes, one in 7 accessions with mitogenomes of about 379 Kb and the other in the longest mitogenome of 712.

### 3.2. Variability of pea mitochondrial genomes

To estimte variability, each of the assembled mitochondrial genomes was compared to that of WL 1238 taken as reference and the observed differences were registered (Appendix B) which are referred to as mutations, including nucleotide substitutions, indels and small inversions (up to 10 bp). The number of different mutations was counted for each gene and intergenic region. The mean value of the 100-fold number of different mutations in the mitochondrial genomes as comparted to that of WL 1238 (Appendix B) per length unit was 0.131 ± 0.03 for coding regions and 0.935 ± 0.069 for non-coding regions. In plastid genomes of the same set of accessions these values comprised 0.386 ± 0.074 for coding regions and 1.955 ± 0.148 for non-coding regions. The proportion of indels among all mutations was about 43% in mitochondrial genomes and about 25% in plastid genomes.

### 3.3. Phylogenetic reconstructions based on updated set of plastid genomes and nuclear His5 gene

In this study we made phylogenetic reconstruction by the Bayesian MCMC method (Fig. 2) based on an almost twice as large (38 vs 22) set of pea plastid genomes than in our previous study (Bogdanova et al., 2018). Expectedly, the major pattern remained the same.

**Figure 2.**
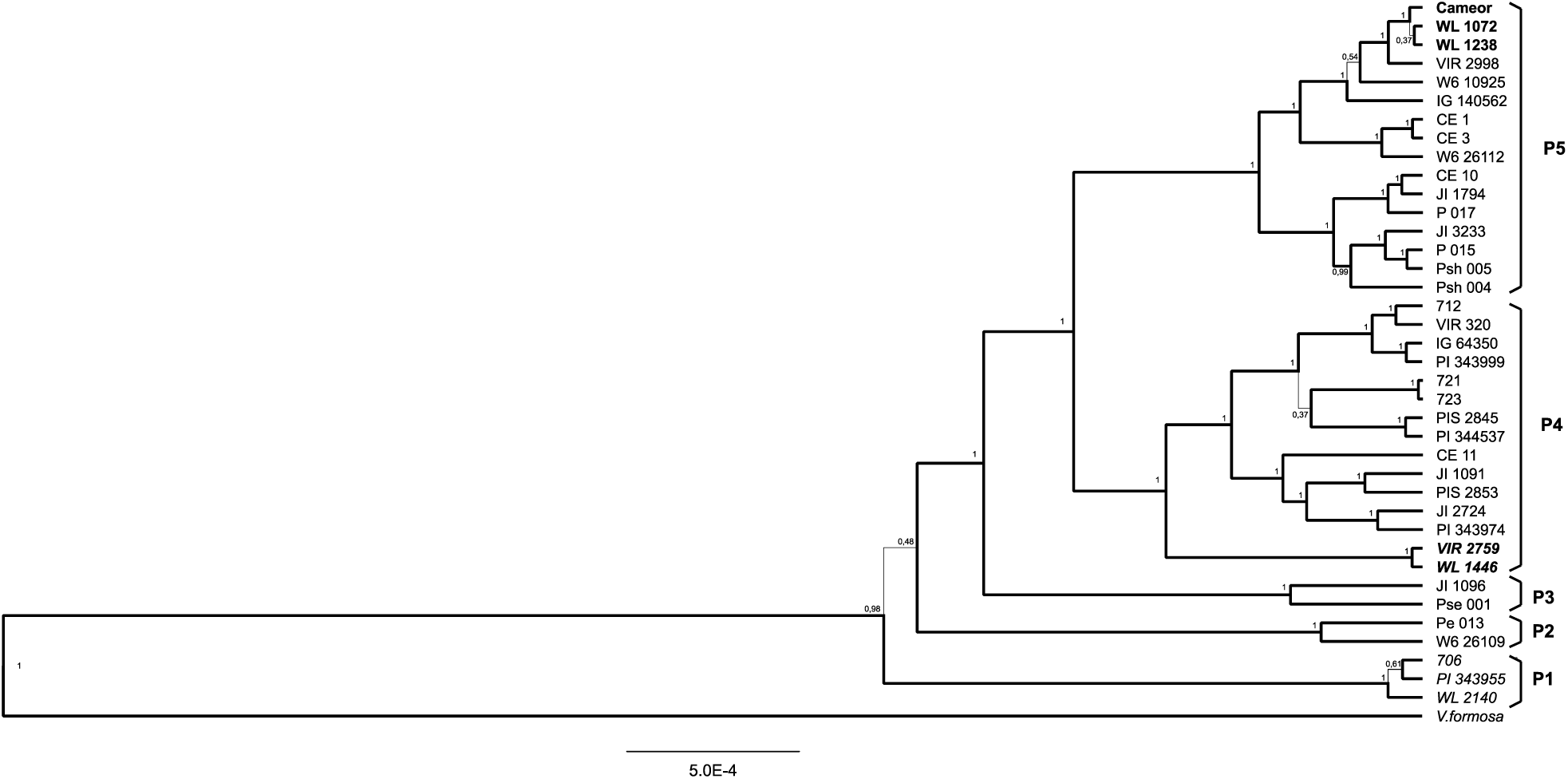
Bayesian reconstruction of phylogeny of the studied pea accessions on the basis of the plastid genomes. Posterior probabilities of the nodes are indicated. Branches corresponding to the nodes supported by the posterior probabilities higher than 0.5 are given as thick lines. Scale bar corresponds to the expected number of nucleotide substitutions per site. Main branches of the tree are designated as P1-P5. Accessions of *Pisum fulvum* are given in italics, of *P. abyssinicum* in bold italic, of the wild subspecies *P. sativum* subsp. *elatius* s.l. in regular Roman, of the cultivated subspecies *P. sativum* subsp. *sativum* in bold. *Vavilovia formosa* serves as outgroup.

For the purpose of comparison with other phylogenetic reconstructions, here we introduce designations of the main revealed branches, being an ordinal number preceded with the letter ‘P’ (for ‘plastid’) (Fig 2).

The most ancient diverged branch P1 contains all the three involved accessions of *P. fulvum* and is opposed to the rest of the tree formed by representatives of *P. sativum*.

The next diverged branch P2 includes wild pea (*P. sativum* subsp. *elatius* s.l.) accessions Pe 013 (Turkey) and W6 26109 (Georgia) and is an ancient plastid lineage “ascending to a ‘missing link’ between *P. fulvum* and *P. sativum*” revelaled eariler (Bogdanova et al., 2018). Its position on the tree is not well resolved, so that we may speak of three main clades of pea plastid genomes of uncertain mutual phylogenetic relationship, P1, P2 and the main *P. sativum* clade. The latter consists of three main branches.

The earliest branch P3 consists of two wild pea (*P. sativum* subsp. *elatius* s.l.) accessions, JI 1096 from Greece and Pse 001 from Turkey. This branch was unexpectedly revealed in our earlier study being represented by the only accession JI 1096 (Bogdanova et al., 2018), here this branch has gained additional support by the second pea accession.

The large branch P4 includes diverse wild pea (*P. sativum* subsp. *elatius* s.l.) accessions and corresponds to the ‘lineage AC’ in our earlier terminology (Zaytseva et al., 2015; Bogdanova et al., 2018). The earliest divergence inside this branch is formed by the accessions (VIR 2759 and WL 1446) of cultivated *P. abyssinicum*.

The third branch P5, also very large, includes accessions lacking *Hsp* AI recognition site in the *rbcL* allele (marker combination B) and corresponds to the ‘lineage B’. Most of them are wild peas, *P. sativum* subsp. *elatius* s.l.. It further branches to form a subbranch of seven accessions with about 3.5-Kb inversion in the plastid genome (Palmer et al., 1985; Bogdanova et al., 2015) and a subbranch of nine accessions without such inversion, including cultivated peas. The three involved representatives of the common cultivated subspecies *P. sativum* subsp. *sativum* (Cameor, WL 1238 and WL 1072) are tightly clustered inside the latter subbranch. They form a larger cluster together with three *P. sativum* subsp. *elatius* accessions from Georgia (VIR 2998), Bulgaria (W6 10925) and Azerbaijan (IG 140562), which are wild peas yet sharing with cultivated peas the specific deletion in the *trnH-psbA* intergenic spacer (Zaytseva et al., 2017). Indeed, the plastid genomes of these three accessions are closely related to those of the cultivated peas, but they are not interspersed with them on the tree and occupy positions outside the ‘cultivated cluster’.

The phylogenetic tree based on the updated set of sequences of the nuclear *His5* gene has been reconstructed (Fig. 3). Its topology is in general similar to that of the tree based on the complete plastid genomes (Fig. 2) but is expectedly less resolved. Like in our earlier reconstruction (Bogdanova et al., 2018), four main clades are resolved: (i) of *P. fulvum*; (ii) of accession Pe 013; (iii) ‘lineage AC’ and (iv) ‘lineage B’.

**Figure 3.**
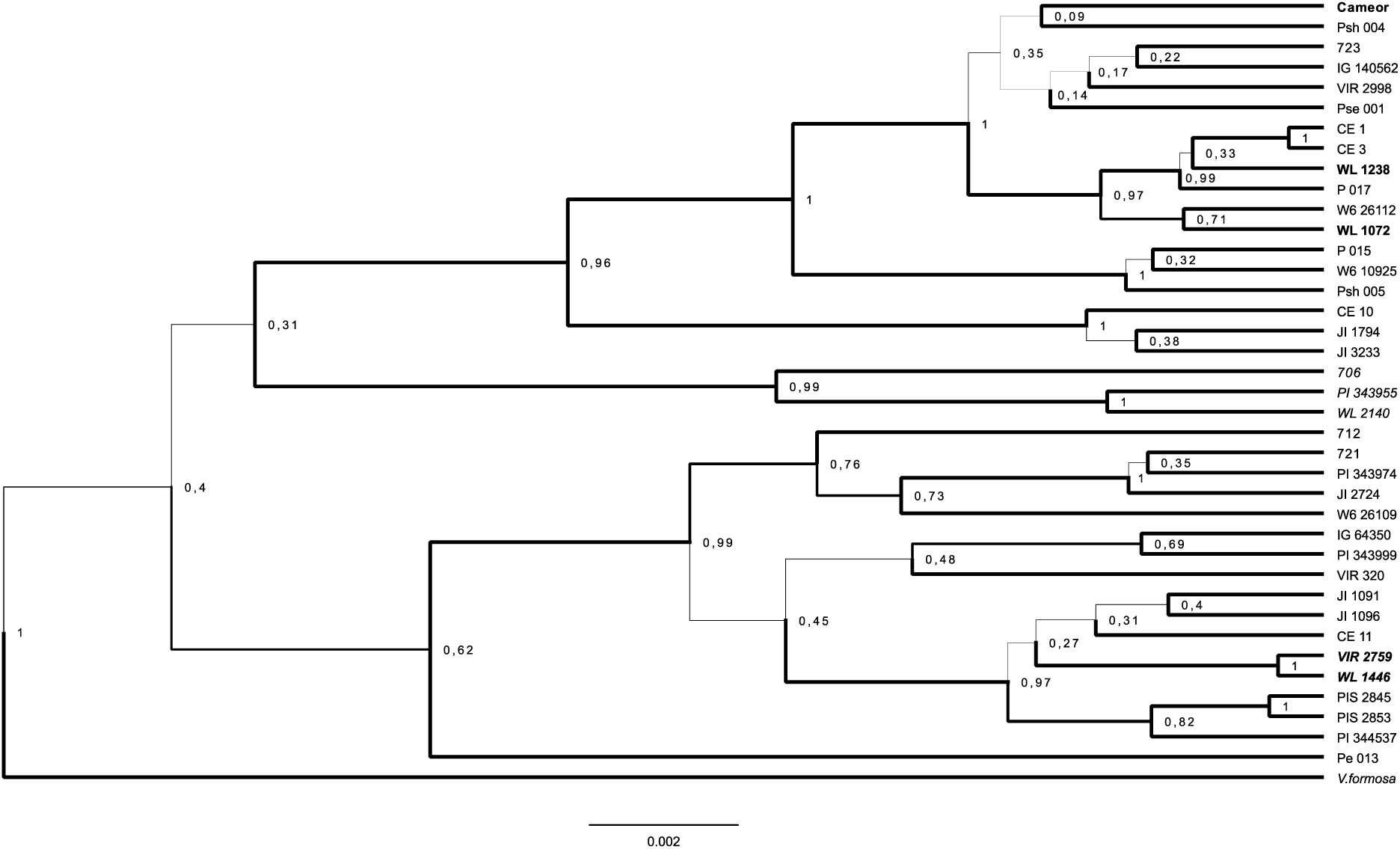
Bayesian reconstruction of phylogeny of the studied pea accessions on the basis of the nuclear *His5* gene. *Vavilovia formosa* serves as outgroup. Designation of taxonomic attribution as in Fig. 2.

The differences from the plastid topology concern positions of two accessions. First, accession W6 26109 (Georgia) in the nuclear gene tree is found inside the ‘lineage AC’ while in the plastid tree it is clustered with Pe 013 to form the ancient plastid lineage P2. The corresponding ancient lineage is also detected in the *His5* tree but it is represented solely by Pe 013.

Second, accession 723 with combination A appeared inside ‘lineage B’. This case is explainable since this accession, from Sardinia, although classified as ‘*Pisum elatius*’ by Ben-Ze’ev and Zohary (1973), has non-dehiscing pods and obviously is a result of hybridistion of wild and cultivated peas, as mentioned in Table 1.

### 3.4. Phylogenetic reconstructions based on mitochondrial genomes

The phylogenetic tree reconstructed by the Bayesian MCMC method from sequences of the mitochondrial genomes (Fig. 4) differed from the above considered trees drastically. Its six main unusual features were as follows:

**Figure 4.**
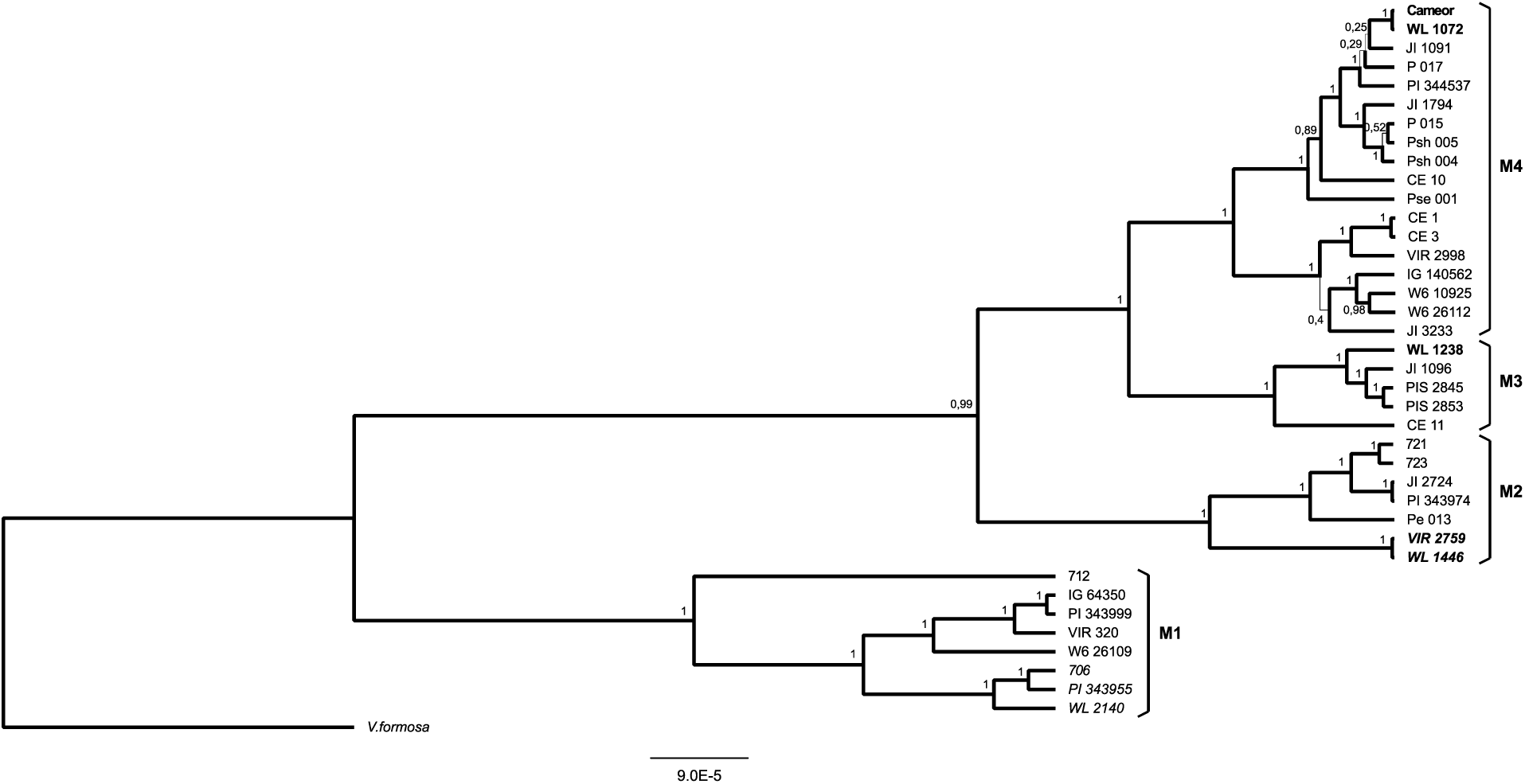
Bayesian reconstruction of phylogeny of the studied pea accessions on the basis of the mitochondrial genomes. *Vavilovia formosa* serves as outgroup. Main branches of the tree are designated as M1-M4. Designation of taxonomic attribution as in Fig. 2.

i. It contains three main clades of unresolved phylogenetic relationship, one of them represented by *V. formosa*, assumed to be the outgroup, and two others by peas, that is, the genus *Pisum* appears on this tree subdivided into two clades of the same rank as the genus *Vavilovia.*
ii. The species *P. fulvum* is not opposed to *P. sativum*. One of the above mentioned principal *Pisum* clades includes *P. fulvum* and some representatives of *P. sativum* subsp. *elatius* s.l. of the lineage AC, the *P. fulvum* branch being immersed inside this clade. The other clade embraces the rest of *P. sativum*, wild and cultivated, and the accessions of *P. abyssinicum*.
iii. The mitochondrial genomes of the accessions forming the ancient branch P2 in the plastid tree (Fig. 2), nominated to be a ‘missing link’ between *P. fulvum* and *P. sativum*, have no unusual characteristics. In the mitochondrial tree (Fig. 4) they are interspersed with representatives of the lineage AC.
iv. Accessions of ‘lineage AC’ and ‘lineage B’ do not form monophyletic clades in the mitochondrial tree, while do so in the plastidic and *His5* trees.
v. There is a well resolved cluster of accessions with mitochondrial genomes lacking the *Psi* I restriction site in the *cox1* gene, that is separating carriers of the combinations B and C from carriers of the ancestral combination A possessing the *Psi* I recognitions site.
vi. While two of the three representatives of the common cultivated subspecies *P. sativum* subsp. *sativum* are tightly clustered, the third, WL 1238, is found in another big cluster.

To distinguish the main branches on the phylogenetic trees based on the mitochondrial genomes (Fig. 4) we use designations starting with the letter ‘M’ (‘mitochondria’).

The branch M1 includes all the three accessions of *P. fulvum* and five representatives of *P. sativum* subsp. *elatius* s.l. with marker combination A, from Algeria, Israel, Turkey and Georgia. Among them we find VIR 320 and 712 known to have plastids incompatible with nuclear genome of cultivated pea (Bogdanova et al., 2015), and W6 26109 – a Georgian accession from the ancient plastid lineage P2 (Bogdanova et al., 2018).

The branch M2 embraces accessions of *P. sativum* subsp. *elatius* s.l. (including Pe 013 from the ancient plastidic lineage P2) and the two accessions of *P. abyssinicum*, all with marker combination A.

The branch M3 includes four accessions of *P. sativum* subsp. *elatius* s.l. from Southern Europe with combination C and the testerline WL 1238 representing the cultivated subspecies.

The branch M4 includes the rest *P. sativum* subsp. *elatius* s.l. with combinations B and C and two representatives of the cultivated subspecies *P. sativum* subsp. *sativum* (Cameor and WL 1072).

## 4. Discussion

### 4.1. Variation of pea mitogenomes

It is well known that plant mitochondrial genomes exist as several structural isoforms (circular, linear or branched) permanently undergoing rearrangements at their repeated segments (Gualberto et al., 2014; Kozik et al., 2019). The here assembled mitogenomes represent isoforms, where possible, consisting of one master circle,. A structural type consisting of two master circles was revealed for the mitogenome of a close pea relative *V. formosa* (Shatskaya et al., 2019), while that in the genus *Silene* L. is represented by up to 128 circular ‘chromosomes’ (Sloan et al., 2012). A substantial structural variation in mitochondrial genomes was found e.g. in the genus *Gossypium* L. (cotton) (Chen et al., 2017) and family Oleaceae (Van de Paer et al., 2017). The enhanced rearrangement processes in the mitochondrial genomes as compared to the plastid genomes is the general trend in plants (Smith and Keeling, 2015). Existence in peas of six different structural types of mitogenomes is hence not surprising.

Of the six revealed structural types of mitochondrial genomes, one type embraced all accesssions of the lineages B and C (those lacking *Psi* I recognitionn site in the *cox1* gene), while mitogenomes of the lineage A were more structurally variable and possessed one of the rest five structural types revealed. Two of the structural types were confined to the M1 clade and three – to the M2 clade. For the purpose of alignment non-collinear mitogenomes were manually cut and the fragments then put in the order and orientation corresponding to that of WL 1238. The number of pieces into which a given mitogenome should be cut to form a collinear reconstruction can serve as a measure of rearrangement events. This number was estimated as five for mitogenomes of the M2 clade, except for two abyssinian peas, which were separated by ten (VIR 2759) and 12 (WL 1446) rearrangements. Mitogenomes of the M1 clade differed from that of WL 1238 by 13 rearrangements except for 712 which differed by 10 rearrangements. Notably, the mitogenome structural types differed in the two accessions representing the same taxon *P. abyssinicum* which is rather uniform morphologically but variable karyologically (Kosterin, 2017).

Chen et al. (2017) reported that in spite of a very high rate in rearrangements of the mitochondrial genome in cotton, the rate of their molecular evolution in terms of nucleotide substitutions was only ca 1/6 of that of the plastid genomes. The similar difference can be noted comparing the here made reconstructions for plastid (Fig. 2) and mitochondrial (Fig. 4) genomes. The given scale bars, although of the same physical size, correspond to 9×10^−5^ nucleotide substitution per site for the mitochondrial tree and 5×10^−4^ for the plastid tree, the former being ca 1/5 of the latter and is very close to the estimate by Chen et al. (2017). Our estimates of the number of different nucleotide substitutions per length unit (including coding and non-coding regions) provided 3.17 ± 0.37 ×10^−3^ for mitochondrial genomes and 8.49 ± 0.67 ×10^−3^ for plastid genomes.

### 4.2. Discordant evolution of oranelles in peas

The phylogenetic reconstruction based on the mitochondrial genomes exhibits multiple discordances from that based the plastid genomes, as illustrated in Fig. 5. The most remarkable feature is that in the mitochondrial tree *P. fulvum* is no more the first branch of the genus but is clustered with five representatives of *P. sativum* subsp. *elatius*. Accessions Pe 013 and W6 26109 do not form any ‘ancient lineage’ as in the plastid genome tree. Accessions lacking the *Hsp* AI restriction site in the plastid *rbcL* gene form a monophyletic clade in the plastid tree and the accessions lacking the *Psi* I restriction site in the mitochondrial *cox1* gene form a monophyletic clade in the mitochondrial tree, but not vice versa. The last but not least, three representatives of the cultivated subspecies *P. sativum* subsp. *sativum* lost monophyly in the mitochondrial genome tree. These obvious discordances can only be interpreted through multiple hybridisation events between divergent pea lines which took place in the past and resulted in discordant inheritance of plastids and mitochondria. Signs of reticulate evolution in the genus *Pisum* were earlier traced, with the use of the Structure software, in phylogeny of a set of nuclear genes by Jing et al. (2007). Now we get its unequivocal evidence from the organellar phylogenies.

**Figure 5.**
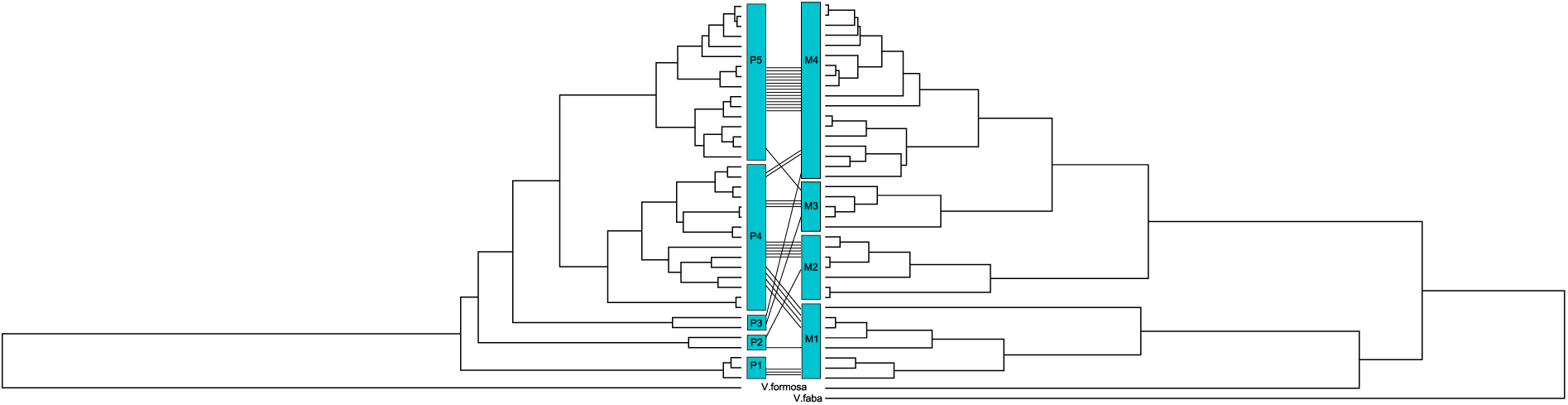
Comparison of phylogenetic trees reconstructed on the basis of plastid (left) and mitochondrial (right) genomes. Designations of main tree branches correspond to those of Fig. 2 and Fig. 4.

Spontaneous hybridisation of wild peas is not surprising. Although pea is often considered to be a selfer, it has conspicuous flowers and occasional cross-pollination by bees does occur even in cultivated peas, especially in traditional local landraces from southern regions (Loening, 1984), since the pistil retains some ovules not yet fertilised with the own pollen when flower opens (Bogdanova and Berdnikov, 2000). Population genetic studies by Smýkal et al. (2018) evaluated the level of cross-pollination in wild peas in SE Turkey as 10–53%. It is geographical isolation of wild peas as well as topographical isolation of their tiny populations from each other, rather than biological inability of crossing, which reduces probability of hybridisation, yet it did occur many times in the course of *Pisum* evolution (Jing et al., 2007; this paper).

In contrast to the substantial discordance of the phylogenies based on mitochondrial and plastid genomes in peas revealed in this study, the similar comparison in the family Oleaceae showed unequivocal concordance in their topologies (Van der Paer et al., 2017).

The nuclear genomes should contain more sophisticated signatures of the hybridisation events which occurred during *Pisum* evolution, and SNP analysis, already attempted for peas (Smýkal et al., 2017; Kreplak et al., 2019), can shed light on them. However, the organellar genomes have an advantage of being free of meiotic recombination and hence are good landmark of evolutionary pathways.

### 4.3. Position of Vavilovia

In the reconstructed mitochondrial genome tree (Fig. 4), one of the pea branches is disposed very close to *Vavilovia* while the rest of peas reside on a long ‘stem’. This is in contrast to the phylogenetic tree based on the plastid genomes where the entire genus *Pisum* crowns a long ‘stem’ sprouting after divergence from the common ancestor with *Vavilovia* (Fig. 2). This could be interpreted in terms of an accelerated evolution of mitochondrial genomes in one particular evolutionary lineage leading to the last common ancestor of the ‘extended’ branch. However, it would difficult to suppose any biological reason for this. More likely *Vavilovia* is not too appropriate outgroup for phylogenetic reconstruction based on the mitochondrial genome.

*Vavilovia* is traditionally considered a genus most close to *Pisum*, yet it differs from peas by perennial life cycle and a non-climbing habitus, being a low plant specialised to highland screes in the Caucasus and Anterior Asia (Vishnyakova, 2020). It looked the most fit outgroup for phylogenetic reconstruction in peas; for this purpose we obtained the sequences of plastid and mitochondrial genomes of *Vavilovia formosa* (Shatskaya et al., 2019). It served a good outgroup in reconstructing phylogeny of peas based on histone H1 genes (Zaytseva et al., 2015; this paper, Fig. 3) and the plastid genomes (Fig. 2), but, as noted above, offered a distorted tree when used for the mitochondrial genome reconstruction (Fig. 4). Addition to the analysis of the mitochondrial genome of a much more remote pea relative *Vicia faba* resulted in a remarkable tree (Fig. 6), in which *Vavilovia* clusters rather tightly with the pea mitochondrial branch M1. This can be interpreted as an ancient introgression of mitochondria from the ancestor of *Vavilovia* to the ancestors of peas united in the branch M1, or vice versa. Remarkably, the present-day *Vavilovia* and *Pisum* are too different plants inhabiting too different habitats to hybridise. Since it is *Vavilovia* which look highly specialised ecologically and morphologically in the entire tribe Fabea, it can be supposed that its ancestor had acquired pea mitochondria still before that specialisation took place and it still resembled peas in its habitus and life strategy.

**Figure 6.**
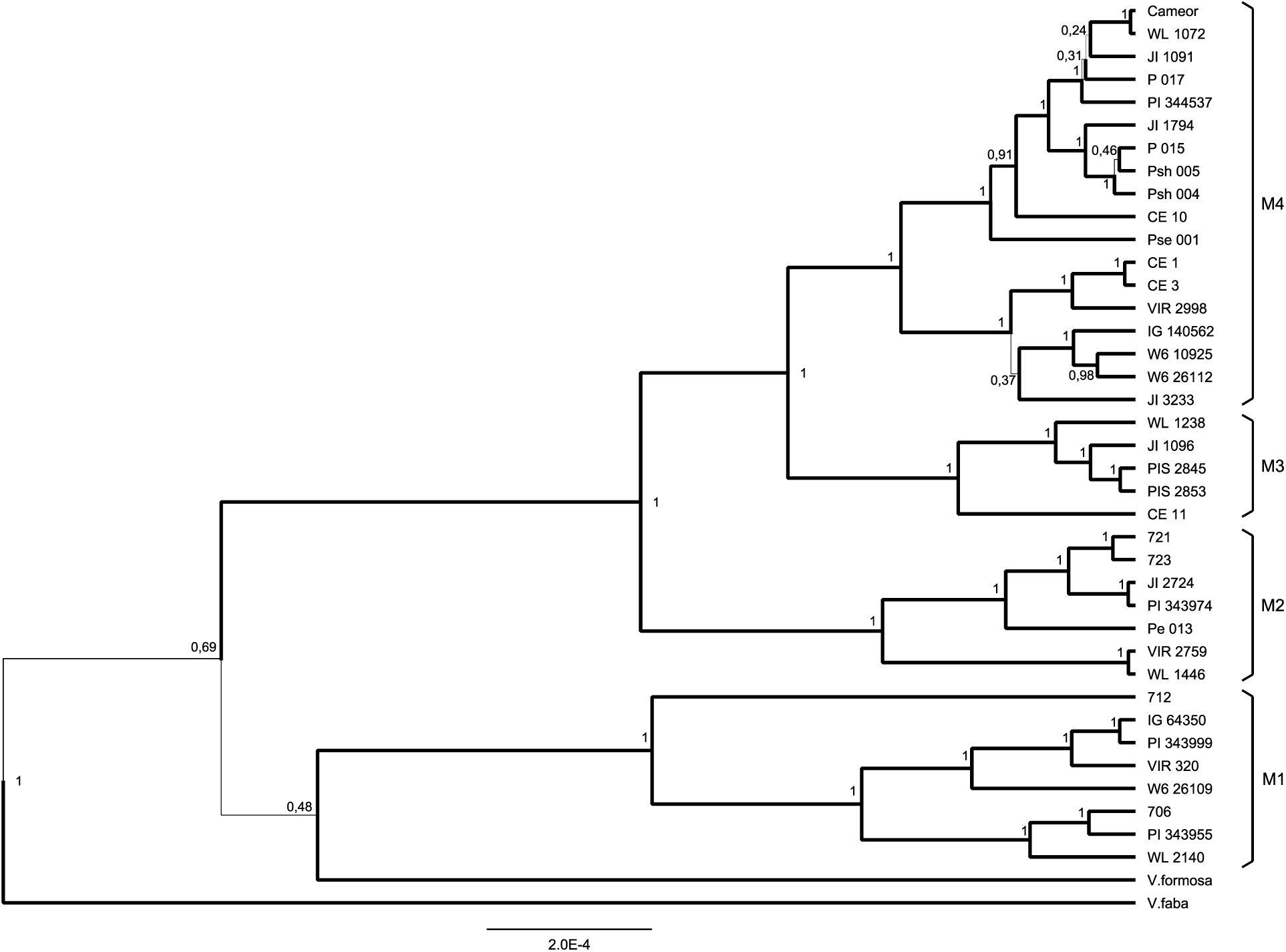
Bayesian reconstruction of phylogeny of the studied pea accessions, *Vavilovia formosa* and *Vicia faba*, on the basis of the mitochondrial genomes. Designations as in Fig. 4.

### 4.4. Organellar constitution and the number of past hybridisation events

Earlier we introduced a system of combinations (A, B, C) of three molecular markers from the three cellular genomes to denote certain presumed lineages of pea evolution (Kosterin and Bogdanova, 2008; Kosterin et al.. 2010; Zaytseva et al., 2017). In view of the here presented new insight in phylogeny of the plastid and mitochondrial genomes, this system becomes insufficiently informative. For instance, it does not reflect belonging to some ‘ancient lineages’. To facilitate discussion below we classify peas with respect to constitution of their organellar genomes as attributed to their main evolutionary branches with were denoted in sections 3.3 and 3.4. For instance, accession Pe 013 has constitution P2 M2, accession W6 26109 - P2 M1, accession 712 - P3 M1, cultivar Cameor – P5 M4, and so on. This organellar genomic constitution is indicated in Table 1. Correspondence of the genomic constitutions to the previously adopted marker combinations A – C is shown in Table 2.

**Table 2.** Correspondence of the genomic constitutions in the sense of this paper to combinations A, B, C of alleles of three molecular markers *rbcL, cox1* and *SCA* from different cellular genomes according to Kosterin and Bogdanova (2008) and Kosterin et al. (2010)/

| combination | organellar genomic constitution | taxon (accessions) |
| --- | --- | --- |
| A | P1 M1 | *P. fulvum* (WL 2140, 706, PI 343955) |
|  | P2 M1 | *P. sativum* subsp. *elatius* (W6 26109) |
|  | P2 M2 | *P. sativum* subsp. *elatius* (Pe 013) |
|  | P4 M1 | *P. sativum* subsp. *elatius* (VIR 320, 712, IG 64350, PI 343999) |
|  | P4 M2 | *P. sativum* subsp. *elatius* (721, 723, JI 2724, PI 343974, Pe 013),<br>*P. abyssinicum* (WL 1446, VIR 2759) |
| C | P3 M3 | *P. sativum* subsp. *elatius* (JI 1906) |
|  | P3 M4 | *P. sativum* subsp. *elatius* (Pse 001) |
|  | P4 M3 | *P. sativum* subsp. *elatius* (CE 11, PIS 2845, PIS 2853) |
|  | P4 M4 | *P. sativum* subsp. *elatius* (JI 1091, PI 344537) |
| B | P5 M3 | *P. sativum* subsp. *sativum* (WL 1238) |
|  | P5 M4 | *P. sativum* subsp. *elatius* (CE 3, CE 1, W6 10925, W6 26112, JI 3233, VIR 2998, P 017, Psh 004, Psh 005, CE 10, P 015, IG 140562, JI 1794)<br>*P. sativum* subsp. *sativum* (WL 1072, Cameor) |

Note: Combination D, differing from combination B by the allele of the nuclear gene *SCA*, was found in some pea accessions with traces of hybridisation during reproduction but not in accessions of genuine wild peas. Accession PI 344537 (Sicily) has combination C and was earlier classified as having combination D (Zaytseva et al., 2017) due to a technical error (adopted from Kosterin and Bogdanova, 2008).

Each case in the above mentioned constitutions where the same plastid branch is combined with more than one mitochondrial branch should be considered as evidence of an independent hybridisation event in the past. We can indicate six such cases: P2 with M1 or M2, P3 with M3 or M4, P4 with M1, M2, M3 or M4, P5 with M3 and M5. This is the lower estimate of the number of hybridisation events with discordant organellar inheritance in the evolution of *Pisum*. There could be more such events resulting in combinations of plastids and mitochondria of different sub-branches of their main evolutionary branches.

### 4.5. Role of nuclear-cytoplasmic incompatibility in shaping organellar phylogeny

The plant phenotype is mostly a product of its tens of thousands nuclear genes. We sequenced only one of them, *His5*, encoding a minor H1 histone subtype and not involved into the genetic program for morphogenesis. Nevertheless, the phylogenetic tree reconstructed on the basis of the nucleotide sequence of this gene (Fig. 3) mostly agrees with expectations from the point of view of morphology and taxonomy; that is evolution of a single gene chosen for its high variability (Zaytseva et al., 2012) appears to have been in line with the mainstream pea evolution, more or less reflected in its taxonomy. The principal topology of the phylogenetic trees reconstructed by Kreplak et al. (2019: Fig. 6) on the basis of 2 thousand nuclear SNP is the same as in our tree based on a single nuclear gene (Fig. 3): the first branch of *P. fulvum* and further bifurcation into a branch with a part of accessions of *P. sativum* ssp. *elatius* s.l. (including ‘*P. humile*’) plus *P. abyssinicum* and a branch with the rest of *P. sativum* subsp. *elatius* plus *P. sativum* subsp. *sativum*. Unfortunately the analysis by Kreplak et al. (2019) included only seven accessions of *P. sativum* ssp. *elatius* s.l. (with only one of them, 721, in common with our analysis), although this inclusive subspecies embraces most of the species’ diversity.

The tree reconstructed on the basis of the plastid genomes has two remarkable discordances with taxonomy and the nuclear gene tree, while the mitochondrial tree looks randomly reshuffled as compared to both of them. This evidences for occasional hybridisation to have taken place in the course of pea evolution, which generated new combinations of nuclear genes and organellar genomes. Two circumstances are important:

i. the plastid and mitochondrial genomes had been discordantly inherited by the hybrids, that would be impossible with strict maternal inheritance of the organelles and,
ii. the plastids were more often inherited concordantly with the sequenced nuclear gene, as well as unknown nuclear genes controlling the external phenotype, while inheritance of the mitochondria was more often discordant.

These facts can be explained through phenomena discovered by our team in remote pea crosses. Biparental and even paternal inheritance of plastids does occur in peas in the situation of nuclear-plastid conflict (Bogdanova and Kosterin 2006; Bogdanova, 2007; Bogdanova et al.; 2014; Kosterin and Bogdanova 2015) resulting from incompatibility of the plastid-encoded subunit of the plastidic heteromeric acetyl-coA carboxylase with one or two its subunits encoded in the nucleus (Bogdanova et al., 2009, 2012; 2015). Milder forms of this conflict appeared widespread in remote crosses involving different wild peas (Bogdanova et al., 2014), to make some combinations of certain plastids and some nuclear genes unviable or, more often, to result in complete or partial loss of fertility. It is noteworthy that the conflict can be overcome by occasional paternal plastid inheritance, so that the presence of paternal plastids in some plant sectors restores their normal phenotype (Bogdanova and Kosterin, 2006; Bogdanova, 2007). In evolutionary terms this means that not any combination of plastids with certain nuclear genes is allowed to be inherited by descendants of occasional spontaneous hybrids, that is, plastid inheritance is to some extent controlled by the nucleus.

At the same time, neither a conflict between the nucleus and mitochondria nor biparental or paternal inheritance of the latter was so far found in peas (Bogdanova, 2007). This means that the type of mitochondria inherited by descendants of a hybrid is determined only by the direction of the cross and is invariably maternal, and mitochondrial evolutionary lineages were not controlled by the nucleus. In some sense the mitochondrial tree shows the history of crosses between diverged peas in the course of the *Pisum* evolution.

As discussed above, comparison of the phylogenetic trees reconstructed from the plastid and mitochondrial genomes suggested independent cases of hybridisation between divergent evolutionary lineages to take place in the *Pisum* evolutionary history. They result in a number of organellar constitutions enumerated above, which, in our earlier system, are manifested in the marker combinations A, B and C. It is remarkable that certain constitutions, e.g. P5 M1 or P5 M2 (which would not fit these combinations) seem not to occur in wild peas. Earlier Kosterin and Bogdanova (2008) reported four pea accessions (of 89 studied) with unusual combinations of cellular genomes. Two of them, VIR 320* and WL 2123 were highly heterogeneous accessions represented by many genotypes with ‘wild’ and ‘cultivated’ traits found in different combinations and obviously represented segregated progeny of occasional crosses of wild and cultivated peas during reproduction. Accessions P 008 (Turkey) and L 90 (without provenance) were received as wild peas but appeared to have non-dehiscing pods hence the material we tested did not represent genuine wild peas.

We may conclude that in wild pea evolution, natural hybridisation of some peas with marker combinations A and B resulted in peas with plastids from the former and mitochondria from the latter but not vice versa. Since mitochondria are maternally inherited in peas, those with combination C most probably resulted from cases when peas with combination B were spontaneously cross-pollinated with those with combination A. Potential nuclear-plastid incompatibilities occurring in such crosses could have been alleviated by paternal plastid inheritance.

### 4.6. Phylogeographic implications

All phylogenetic reconstructions obtained in this and our earlier (Zaytseva et al;, 2012; 2015) studies revealed a robust monophyletic branch embracing wild and cultivated accessions earlier recognised as ‘lineage B’ marked by the loss of certain restriction sites in the plastid *rbcL* and mitochondrial *cox1* gene, earlier chosen to conventionally classify wild pea genetic diversity (Kosterin, Bogdanova, 2008; Kosterin et al., 2010). As follows from the here presented phylogenetic reconstructions based on the plastid and mitochondrial genomes, each of these two losses obviously occurred once in evolution, *Hsp* AI site in the plastidic *rbcL* was lost at the origins of the plastid branch P5 (Fig. 2) and the *Psi* I site in mitochondrial *cox1* was lost during evolution of ‘stem’ leading to mitochondrial branches M4+M3 (Fig. 4).

Earlier Kosterin et al. (2010) supposed that peas with combination C represented a transitory stage of evolution leading to the ‘lineage B’, when the mitochondrial *cox1* already had lost the restriction site while the plastid *rbcL* not yet, and that the difference between combinations C and B concerned the latter mutation only. In such a scenario, all phylogenetic reconstructions would show a monophyletic branch composed of accessions with combination C and the inner monophyletic branch of ‘lineage B’. The revealed pattern is nothing like this and carriers of combination C do not represent a monophyletic group in any reconstructed tree. In the plastid tree (Fig. 2), the carriers of combination A (with mitochondria of the branches M1 and M2) and C (with mitochondria of the branches M3 and M4) are interspersed in the branch P4 without phylogenetic regularity. Also in the mitochondrial tree (Fig. 4), the carriers of combination B (with plastids of the branch P5) and C (with plastids of the branches P3 and P4) are interspersed without phylogenetic regularity in the branches M3 and M4.

This finding disproves the phylogeographic scenario proposed by us earlier (Kosterin et al., 2010), implying the evolution from carriers of the ancestral combination A of south-eastern Mediterranean to ‘lineage B’ inhabiting south-eastern Mediterranean and Maghreb through ‘intermediate peas’ with combination C inhabiting Southern Europe. The phylogenetic patterns observed can only be interpreted if to assume that ‘combination C’, embracing four different organellar constitutions (Table 2), had not occurred via a loss of *Psi* I restriction site from the *cox1* gene in some representative of the lineage A, but originated from not less than four independent hybridisation events between the representatives of the plastid branches P3 and P4 and the mitochondrial branches M3 and M4.

An important inference from the above is that most of Southern Europe, from Sicily to Hungary and from Portugal to Greece, is inhabited by wild peas (with combination C) of hybrid origin. There we found the organellar constitutions P3 M3 (JI 1096, Greece), P4 M3 (3 accessions, from Portugal, Hungary and Sicily) and P4 M4 (JI 1091, Greece, and PI 344537, Sicily). Curiously, the constitutions P3 M3 and P4 M4 were found in the same series of wild peas collected by Howard Scott Gentry in 1969 at the Karyes Town on the Athos Peninsula, Greece, and constitutions P4 M3 and P4 M4 occur in the two accessions from Sicily.

These presumable hybridisations could take place when wild *P. sativum* started to expand to Europe from the eastern Mediterranean from where the genus *Pisum* had originated. As a result of not less than three such events, the hybrid descendants got the allowed organellar constitution (manifested as marker combination C; one more hybrid organellar constitution corresponding to combination C, P3 M4, was not found in Europe). It should be stressed that they occurred between wild peas and long ago, without participation of cultivated peas (*P. sativum* subsp. *sativum*), since the mitochondrial genome of these European wild peas with combination C do not contain markers of the cultivated pea lineage, e.g. the 7-bp deletion in the *trnH-psbA* spacer (Zaytseva et al., 2017).

Wild peas of ancient hybrid origin with the organellar constitution P3 M4, are also found in Turkey (Pse 001).

The ‘lineage B’ (constitution P5 M4) can be supposed to originate in the Caucasus and Asia Minor, which it currently inhabits (Kosterin et al., 2010; Zaytseva et al., 2017). At present, wild representatives of *P. sativum* carrying plastids and mitochondria from different evolutionary clades, including their hybrid combination P4 M4 (combination C), grow side by side in Turkey, Israel and the Caucasus (Kosterin et al., 2008; Zaytseva et al., 2017; this study).

The accession WL 1238 is assumed to belong to the cultivated subspecies *P. sativum* subsp. *sativum*, yet it was found to possess an unexpected and unusual organellar constitution P5 M3 and hence falls out of ‘lineage B’ (althouth has the marker combination B). In should be noticed that this is a testerline created by Hebert Lamprecht as a multiple hybrid. We failed to trace in its available history (Anonymus, 1984) the source of M3 mitochondria which could be wild pea from Southern Europe, but have to suppose that such ancestor should have existed.

### 4.7. Phylogeny and phenotype

Accessions involved into our analysis of wild representatives of *P. sativum*, conventionally denoted as *P. sativum* subsp. *elatius* in a broad sense, were grown in greenhouse and demonstrated a great diversity with respect to external characters. Many of them possess remarkable and even unique characters. They could motivate erection of a number of taxa below the species rank but fortunately escaped attention of taxonomic botanists. For example, most leaves of Psh 004 (a low pea from E Turkey) are impairipinnate, composed of 3 to 5 leaflets, with the terminal tendril domain replaced with an apical leaflet. Accession PI 343999 (a low pea with strongly dentate leaflets from S Turkey) has a thick, fleshy pod walls (phenotype n), which was hitherto found among wild peas only in *P. fulvum* and was one of its diagnostic characters; at the same time the pods bear neoplastic pustules (phenotype Np) never occurring in *P. fulvum*. Accessions 712 and IG 64350 (low peas with dentate leaflets from S Israel and Algeria, respectively) are characterised by numerous purple dots on pod walls. Young plants of accession P 017 (a medium-tall accession from S Turkey) demonstrate transitory, lasting for 2-3 days, deep purple-violet anthocyanin hue of the foliage of the first 2-3 nodes, while mature plants bear some neoplastic pustules (phenotype Np) on the stem, while normally they appear (in the greenhouse conditions) only on the pod walls. Accession Pse 001 (a tall pea from E Turkey) has a curious warm, carmine-red hue of its large, well pigmented flowers. Accession PI 343974 from SW Turkey is exceptionally tall and has very long pods. Some obviously unrelated accessions of distant provenance have extremely short, almost reduced peduncles, e.g. CE 11 (NE Portugal), JI 1091, JI 1096 (Greece), JI 3233 (Syria); based on this character the latter accession was attributed to *P. sativum* subsp. *elatius* var. *brevipedunculata* Davis thought to occur in Turkey and Cyprus (Davis. 1969). Iranian accessions CE 9 and CE 10 are characterised by extremely fast seed germination (like *P. abyssinicum*). Neither of these morphological peculiarities finds any reflection in the phylogenies reconstructed, as their carriers did not form distinct, early evolutionary branches. At the same time, an unusual character, amphycarpy, was found earlier in one (Pe 013) of the two accessions of the basal evolutionary branch P2 in the phylogenetic tree reconstructed from the plastid genomes (Bogdanova et al., 2018).

## Declaration of conflict of interest

none

## Funding

This work is supported by Russian State Scientific project № 0324-2019-0039-C-01 at the Institute of Cytology & Genetics SB RAS, Novosibirsk and the project № 19-04-00162 of the Russian Fund for Fundamental Research. We thank Clarice Coyne, Michael Ambrose, Norman Weeden and Petr Smykal for providing valuable wild pea germplasm.

High-throughput sequencing was performed at IC&G Center of Genomic Investigations, Sanger sequencing was performed at the SB RAS Genomics Core Facility, genome assembly and phylogenetic reconstruction were carried out with the use of Computational Facility of the Siberian Supercomputer Center SB RAS and Computational Facility of Novosibirsk State University. Plants were grown in the greenhouse of the SB RAS Artificial Plant Growing Facility.

## Appendix A. Supplementary material

Differences registered in the sequenced plastid genomes as compared to that of accession WL1238 (*Pisum sativum* subsp. *sativum*)

## Appendix B. Supplementary material

Differences registered in the sequenced mitochondrial genomes as compared to that of accession WL1238 (*Pisum sativum* subsp. *sativum*)

## Notes

### Competing Interest Statement

The authors have declared no competing interest.

## References

Abbo, S., Zesak, I., Schwartz, E., Lev-Yadun, S., Gopher, A. 2008. Experimental harvesting of wild peas in Israel: implications for the origins of Near East farming. Journal of Archaeological Science 35: 922–929. https://doi.org/10.1016/j.jas.2007.06.016.

Abbo, S., Lev-Yadun, S., Heun, M. Gopher, A. 2013. On the ‘lost crops’ of the neolithic Near East. Journal of Experimental Botany 64: 815–822. https://doi.org/10.1093/jxb/ers373.

Altschul S.F., Gish W., Miller W., Myers E.W., Lipman D.J. 1990. Basic local alignment search tool. Journal of Molecular Biology. 215: 403–410. https://doi.org/10.1016/S0022-2836(05)80360-2

Ambrose, M.J., Ellis, T.H.N. 2008. Ballistic seed dispersal and associated seed shadow in wild *Pisum* germplasm. Pisum Genetics. 40, 5–10.

Anonymus. 1984. The Pisum-Genebank. Listings of Weibullsholm Collection. Nordic Gene Bank, Weibullsholm Plant Breeding Institute, p. 1–345.

Ben-Ze’ev, N., Zohary, D. 1973. Species relationship in the genus *Pisum* L. Israel J. Bot. 22, 73–91.

Bogdanova, V.S. 2007. Inheritance of organelle DNA markers in a pea cross associated with nuclear-cytoplasmic incompatibility. Theor. Appl. Genet. 114, 333–339. https://doi.org/10.1007/s00122-006-0436-6

Bogdanova, V.S., Berdnikov, V.A. 2000. A study of potential ability for cross-pollination in pea originating from different parts of the world. Pisum Genetics. 32, 16–17.

Bogdanova, V.S., Galieva, E.R., Kosterin, O.E. 2009. Genetic analysis of nuclear-cytoplasmic incompatibility in pea associated with cytoplasm of an accession of wild subspecies *Pisum sativum* subsp. *elatius* (Bieb.) Schmahl. Theor. Appl. Genet. 118, 801–809. https://doi.org/10.1007/s00122-008-0940-y

Bogdanova, V.S., Galieva, E.R., Yadrikhinskiy, A.K., Kosterin, O.E. 2012. Inheritance and genetic mapping of two nuclear genes involved in nuclear-cytoplasmic incompatibility in peas (*Pisum sativum* L.). Theor. Appl. Genet. 124, 1503–1512. https://doi.org/10.1007/s00122-012-1804-z

Bogdanova, V.S., Mglinets, A.V., Shatskaya, N.V., Kosterin, O.E., Solovyev, V.I., Vasiliev, G.V. 2018. Cryptic divergences in the genus *Pisum* L. (peas), as revealed by phylogenetic analysis of plastid genomes. Mol. Phyl. Evol. 129, 280–290. https://doi.org/10.1016/j.ympev.2018.09.002

Bogdanova, V.S., Kosterin, O.E. 2006. A case of anomalous inheritance of chloroplasts in crosses of the garden pea with participation of one of the wild forms. Doklady Akademii Nauk. 406, 256-259 (in Russian).

Bogdanova, V.S., Kosterin, O.E., Yadrikhinskiy, A.K. 2014. Wild peas vary in their cross-compatibility with cultivated pea (*Pisum sativum* subsp. *sativum* L.) depending on alleles of a nuclear-cytoplasmic incompatibility locus. Theor. Appl. Genet. 127, 1163–1172. https://doi.org/10.1007/s00122-014-2288-

Bogdanova, V.S., Zaytseva, O.O., Mglinets, A.V., Shatskaya, N.V., Kosterin, O.E., Vasiliev, G.V. 2015 Nuclear-cytoplasmic conflict in pea (*Pisum sativum* L.) is associated with nuclear and plastidic candidate genes encoding Acetyl-CoA carboxylase subunits. PLoS ONE. 10 (3), e0119835. https://doi.org/10.1371/journal.pone.0119835.

Chen, Z., Nie, H., Wang, Y., Pei H., Li, S., Zhang, L., Hua, J. 2017. Rapid evolutionary divergence of diploid and allotetraploid *Gossypium* mitochondrial genomes. BMC Genomics 18: 876 http://doi.org/10.1186/s12864-017-4282-5.

Chevreux, B., Wetter, T., Suhai, S. 1999. Genome sequence assembly using trace signals and additional sequence information. Computer Science and Biology: Proceedings of the German Conference on Bioinformatics (GCB). 99, 45–56.

Coulot, P., Rabaute, P. 2016. Monographie de Leguminosae de France. Tome 4. Tribus des Fabeae, des Cicereae et des Genisteae. Bulletin de la Société Botanique du Centre-Ouest. 46, 1–902.

Coulot, P., Rabaute, P. 2017. Deuxièmes compléments à la Monographie des *Leguminosae* de France. Le Monde des Plantes 516:11–35.

Darriba, D., Taboada, G.L., Doallo, R., Posada, D. 2012. jModelTest 2: more models, new heuristics and parallel computing. Nature Methods. 9 (8), 772. https://doi.org/10.1038/nmeth.2109

Davis, H. 1969. Materials for a flora of Turkey: XIX. Legumiosae: Vicieae. Notes of Royal Botanical Garden of Edinbourgh 29 (3): 311–320.

Drummond, A.J., Rambaut, A. 2007. BEAST: Bayesian evolutionary analysis by sampling trees. BMC Evol. Biol. 7, 1. https://doi.org/10.1186/1471-2148-7-214

Ellis, T.H.N., Poyser, S.J., Knox, M.R., Vershinin, A.V., Ambrose, M.J. 1998. Polymorphism of insertion sites of *Ty1-copia* class retrotransposons and its use for linkage and diversity analysis in pea. Mol. General Genet. 260, 9–19.

Gualberto, J.M., Mileshina, D., Wallet, C., Niazi, A.K., Weber-Lotfi, F., Dietrich, A. 2014. The plant mitochondrial genome: dynamics and maintenance. Biochimie 100: 107–120. https://doi.org/10.1016/j.biochi.2013.09.016.

Guindon, S., Gascuel, O. 2003. A simple, fast and accurate method to estimate large phylogenies by maximum-likelihood. Systematic Biology. 52, 696–704. https://doi.org/10.1080/10635150390235520

Jansen, R.K., Raubeson, L.A., Boore, J.L., dePamphilis, C.W., Chumley, T.W., Haberle, R.C., Wyman, S.K., Alverson, A.J., Peery, R., Herman, S.J., Fourcade, H.M., Kuehl, J.V., McNeal, J.R., Leebens-Mack, J., Cui, L. 2005. Methods for obtaining and analyzing whole chloroplast genome sequences. Methods Enzymol. 395, 348–384. https://doi.org/10.1016/S0076-6879(05)95020-9

Jing, R.C., Knox, M.R., Lee, J.M., Vershinin, A.V., Ambrose, M., Ellis, T.H.N., Flavell, A.J. 2005. Insertional polymorphism and antiquity of PDR1 retrotransposon insertions in *Pisum* species. Genetics. 171, 741–752. https://doi.org/10.1534/genetics.105.045112

Jing, R., R. Johnson, A. Seres, G. Kiss, M. J. Ambrose, Knox, R., Ellis, T.H.N., Flawell, A.J. 2007. Gene-based sequence diversity analysis of wild pea (*Pisum*). Genetics. 177, 2263–2275. https://doi.org/10.1534/genetics.107.081323

Jing, R., Vershinin, A., Grzebota, J., Shaw, P., Smýkal, P., Marshall, D., Ambrose, M.J., Ellis, T.H.N., Flavell, A.J. 2010. The genetic diversity and evolution of field pea (*Pisum*) studied by high throughput retrotransposon based insertion polymorphism (RBIP) marker analysis. BMC Evolutionary Biology. 10, 44. https://doi.org/10.1186/1471-2148-10-44

Kosterin, O.E. 2016. Prospects of the use of wild relatives for pea breeding. Russian Journal of Genetics: Applied Research. 6 (3), 233–243. https://doi.org/10.1134/S2079059716030047

Kosterin, O.E. 2017. Abyssinian pea (Lathyrus schaeferi Kosterin nom. nov. pro *Pisum abyssinicum* A. Br.) is a problematic taxon. Vavilovskii Zhurnal Genetiki i Selektsii = Vavilov Journal of Genetics and Breeding. 1 (2), 158–169. https://doi.org/10.18699/VJ17.234

Kosterin, O.E., Bogdanova, V.S. 2008. Relationship of wild and cultivated forms of *Pisum* L. as inferred from an analysis of three markers, of the plastid, mitochondrial and nuclear genomes. Genet. Res. Crop. Evol. 55, 735–755. https://doi.org/10.1007/s10722-007-9281-y

Kosterin, O.E., Bogdanova, V.S. 2015. Reciprocal compatibility within the genus *Pisum* L. as studied in F1 hybrids: 1. Crosses involving *P. sativum* L. subsp. *sativum.* Genet. Resour. Crop Evol. 62 (5), 691–709. https://doi.org/10.1007/s10722-014-0189-z

Kosterin, O.E., Bogdanova, V.S., Mglinets, A.V. 2020. Wild pea (*Pisum sativum* L. subsp. *elatius* Aschers. et Graebn. s. l.) at the periphery of its range: Zagros Mountains. Vavilovskii Zhurnl Genetiki i Selektsii=Vavilov Journal of Genetics and Breeding 24 (1): 60–68. https://doi.org/10.18699/VJ20.596

Kosterin, O.E., Zaytseva, O.O., Bogdanova, V.S., Ambrose, M. 2010. New data on three molecular markers from different cellular genomes in Mediterranean accessions reveal new insights into phylogeography of *Pisum sativum* L. subsp. *elatuis* (Beib.) Schmahl. Genet. Resour. Crop Evol.. 57, 733–739. https://doi.org/10.1007/s10722-009-9511-6

Kozik, A., Rowan, B.A., Lavelle, D., Berke, L., Schranz, E., Michelmore, R.W., Christensen, A.C.. 2019. The alternative reality of plant mitochondrial DNA: one ring does not rule them all. PLOS Genet. 15 (8): e1008373. https://doi.org/10.1371/journal.pgen.1008373

Kreplak K, Madoui M-A, Cápal P, Novák P, Labadie K, Aubert G, Bayer PE, Gali KK, Syme RA, Main D, Klein A, Bérard A, Vrbová I, Fournier C, d’Agata L, Belser C, Berrabah W, Toegelová H, Milec Z, Vrána J, Lee H, Kougbeadjo A, Térézol M, Huneau C, Turo CJ, Mohellibi N, Neumann P, Falque M, Gallardo K, McGee R, Tar’an B, Bendahmane A, Aury JM, Batley J, Le Paslier MC, Ellis N, Warkentin TD, Coyne CJ, Salse J, Edwards D, Lichtenzveig J, Macas J, Doležel J, Wincker P, Burstin J (2019) A reference genome for pea provides insight into legume genome evolution. Nature Genet. 51: 1411–1422. https://doi.org/10.1038/s41588-019-0480-1

Larkin, M.A., Blackshields, G., Brown, N.P., Chenna, R., McGettigan, P.A., McWilliam, H., Valentin, F., Wallace, I.M., Wilm, A., Lopez, R., Thompson, J.D., Gibson, T.J., Higgins, D.G. 2007. Clustal W and Clustal X version 2.0. Bioinformatics. 23, 2947–2948. https://doi.org/10.1093/bioinformatics/btm404

Loenning, W.-E. 1984. Cross fertilization in peas under different ecological conditions. Pisum Newsletter. 16, 38–40.

Maxted, N. Ambrose, M. 2001. Peas (*Pisum* L.). In: Plant genetic resources of legumes in the Mediterranean. Eds. Maxted N. and S.J. Bennett. Kluwer Academic Publishers. The Netherlands. Chapter 10, pp 181–190.

Maxted, N., Kell, S.P. 2009. Establishment of a global network for the *in situ* conservation of crop wild relatives: status and needs. FAO Commission on Genetic Resources for Food and Agriculture. Rome.

Milne, I., Stephen, G., Bayer, M., Cock, P.J.A., Pritchard, L., Cardle, L., Shaw, P.D., Marshall, D. 2013 Using Tablet for visual exploration of second-generation sequencing data. Briefings in Bioinformatics. 14, 193–202. https://doi.org/10.1093/bib/bbs012

Muehlbauer, F.J., Kaiser, W.J., Kutlu, Z., Sperling, C.R. 1990. Collection of *Pisum* germplasm in Turkey in 1985 and 1989. Pisum Genetics. 22, 98–99.

Palmer, J.D., Jorgensen, R.A., Thompson, W.F. 1985. Chloroplast DNA variation and evolution in *Pisum* : patterns of change and phylogenetic analysis. Genetics. 109, 195-213. PMCID: PMC1202476

Schaefer, H., Hechenleitner, P., Santos-Guerra, A., Menezes de Sequeira, M., Pennington, R.T., Kenicer, G., Carine, M.A. 2012. Systematics, biogeography, and character evolution of the legume tribe Fabeae with special focus on the middle-Atlantic island lineages. BMC Evol Biol. 12, 250. https://doi.org/10.1186/1471-2148-12-250

Shatskaya, N.V., Bogdanova, V.S., Kosterin, O.E., Vasiliev, G.V., Kimeklis, A.K., Andronov, E.E., Provorov, N.A. 2019. The plastid and mitochondrial genomes of Vavilovia formosa (Stev.) Fed. and the phylogeny of related legume genera. Vavilovskii Zhurnal Genetiki i Selektsii = Vavilov Journal of Genetics and Breeding 23 (8), 972–980. https://doi.org/10.18699/VJ19.574.

Siniauskaya, M.G., Makarevich, A.M., Goloenko, I.M., Pankratov, V.S., Liaudanski, A.D., Danilenko, N.G., Lukhanina, N.V., Shimkevich A.M., Davydenko, O.G. 2020. The study of organelle DNA variability in alloplasmic barley lines in the NGS era. Vavilovskii Zhurnal Genetiki i Selektsii = Vavilov Journal of Genetics and Breeding 24 (1), 12–19. https://doi.org/10.18699/VJ19.589

Sloan, D.B., Alverson, A.J., Chuckalovcak, J.P., Wu, M., McCauley, D.E., Palmer, J.D., Taylor, D.R. 2012. Rapid evolution of enormous, multichromosomal genomes in flowering plant mitochondria with exceptionally high mutation rates. PLoS Biology 10 (1): e1001241. https://doi.org/10.1371/journal.pbio.100124.

Smith, D.R., Keeling, P.J. 2015. Mitochondrial and plastid genome architecture: reoccurring themes, but significant differences at the extremes. Proc. Natl. Acad. Sci. USA 112 (33): 10177–10184. https://doi.org/10.1073/pnas.1422049112.

Smýkal, P., Trněný, O., Brus, J., Hanáček, P., Rathore, A., Roma, R.D., Pechanek, V., Douchoslav, M., Battacharrya, D., Bariotakis M., Pirintsos, S., Berger, J., Toker, C. 2018. Genetic structure of wild pea (*Pisum sativum* subsp. *elatius*) populations in the northern part of the Fertile Crescent reflects moderate cross-pollination and strong effect of geographic but not environmental distance. PLoS ONE 13(3), e0194056. https://doi.org/10.1371/journal.pone.0194056

Smýkal, P., Hradilová I., Trněný, O., Brus, J., Rathore, A., Bariotakis, M., Das, R.R., Bhattacharyya, D., Richards, C., Coyne, C.J.. Pirintsos, S. 2017. Genomic diversity and macroecology of the crop wild relatives of domesticated pea. Scientific Reports 7, 17384. https://doi.org/10.1038/s41598-017-17623-4

Tamura, K., Stecher, G., Peterson, D., Kumar, S. 2013. MEGA6: molecular evolutionary genetics analysis version 6.0. Mol. Biol. Evol. 30, 2725–2729. https://doi.org/10.1093/molbev/mst197

Tar’an, B., Zhang, C., Warkentin, T., Tullu, A., Vandenberg, A. 2005. Genetic diversity among varietis and wild species accessions of pea (*Pisum sativum* L.) based on molecular markers, and morphological and physiological characters. Genome. 48, 257–272. https://doi.org/10.1139/g04-114

Van de Paer, C., Bouchez, R., Besnar, G. 2017. Prospect of evolutionary mitogenomics of plants: a case study of the olive family (Oleaceae). Mol. Ecol. Resour. 2017: 1–17. https://doi.org/10.1111/1755-0998.12742

Vershinin, A.V., Allnutt, T.R., Knox, M.R., Ambrose, M.J., Ellis, T.H.N.. 2003. Transposable elements reveal the impact of introgression rather than transposition, in *Pisum* diversity, evolution and domestication. Mol. Biol. Evol. 20, 2067–2075. https://doi.org/10.1093/molbev/msg220

Vishnyakova, M. 2020. The Vavilov’s Institute (VIR) contribution to the survey and study of *Vavilovia formosa* (Fabaceae). Bio. Comm. 65 (1), 28–40. https://doi.org/10.21638/spbu03.2020.103

Weeden, N.F. 2007. Genetic changes accompahying the domestication of *Pisum sativum* : is there a common genetic basis to the ‘domestication syndrome’ for legumes? Ann. Bot. 100, 1017–1025.

Weeden, N. 2018. Domestication of pea (*Pisum sativum* L.): the case of the Abyssinian Pea. Frontiers Plant Sci. 9, 515. https://doi.org/10.3389/fpls.2018.00515

Zaytseva, O.O., Bogdanova, V.S, Kosterin, O.E. 2012. Phylogenetic reconstruction at the species and intraspecies levels in the genus *Pisum* (L.) (peas) using a histone H1 gene. Gene. 504, 192–202. https://doi.org/10.1016/j.gene.2012.05.026

Zaytseva, O.O., Gunbin, K.V., Mglinets, A.V., Kosterin, O.E. 2015. Divergence and population traits in evolution of the genus *Pisum* L. as reconstructed using genes of two histone H1 subtypes showing different phylogenetic resolution. Gene. 556, 235–244. https://doi.org/10.1016/j.gene.2014.11.062

Zaytseva O.O., Bogdanova V.S., Mglinets A.V., Kosterin O.E. 2017. Refinement of the collection of wild peas (*Pisum* L.) and search for the area of pea domestication with a deletion in the plastidic *psbA-trnH* spacer. Genetic Resources and Crop Evolution. 64 (6), 1417–1430. https://doi.org/10.1007/s10722-016-0446-4

Zohary, D., Hopf, M. 2000. Domestication of Plants in the Old World, 3rd ed. Clarendon Press, Oxford.

